# Oxidative stress in two pollinators from different landscapes based on different pesticide residue profiles

**DOI:** 10.1101/2025.10.04.680455

**Authors:** Keiana Briscoe, Mia Byarlay, Reed Johnson, Sandra Rehan, Hongmei Li-Byarlay

## Abstract

Oxidative stress (OX) is a state of imbalance between antioxidants and reactive oxygen species, which are the byproducts of oxidative phosphorylation in the mitochondria. Different landscapes: conventional, organic, and roadside use different pesticides and pest control management practices pest control. The roadside landscape is a control for minimum application of herbicides and management. Two important pollinators, honey bees (*Apis mellifera)* and small carpenter bees (*Ceratina calcarata*) are studied here for their abiotic stress from different landscape. It is unknown whether the differences in these landscapes affect the oxidative stress of these two pollinators. Conventional farms offer the most exposure to pesticides, which have been related to increased stress levels. We hypothesized that honey bees and small carpenter bees (SCB) from conventional farms would experience the highest levels of OX. Results from the lipid assay indicated that honey bees and SCB of organic and roadside landscapes experienced the lowest and highest OX levels. The results of the protein carbonyl assay indicated a significant difference found between conventional and organic bees (p < 0.05), based on Tukey’s post-hoc test. Previous research has shown that feral bees have a tolerance to high stress levels. The same could be true of honey bees and SCB from roadside landscapes, where the conditions imitate natural habitats. Organic farms use naturally derived pesticides and moderate management practices, likely contributing to their lower levels. Examining how these pollinators are affected by different farm landscapes can reveal the benefits and detriments of their corresponding management practices. This will lead to streamlined practices that are healthier for pollinators.

## Introduction

Oxidative stress is a condition of cellular imbalance caused by the accumulation of reactive oxygen species (ROS) (Chaitanya et al. 2016; Li-Byarlay and Cleare 2020). ROS are the byproducts of the necessary metabolic processes that occur in mitochondria (Betteridge 2000). In excess, ROS have been linked to aging, apoptosis, and acute mortality (Nikolenko et al. 2011). In healthy cells, antioxidant compounds remove ROS through reduction or catalysis into water. Under oxidative stress, cells are more susceptible to DNA damage, protein carbonylation, or lipid peroxidation by ROS (El-Demerdash et al. 2018). The effects of oxidative stress have been studied across many species (Kramer et al. 2021; Kujoth et al. 2005; Li et al. 2008; Li-Byarlay and Cleare 2020). Oxidative stress can be induced by a variety of biotic and abiotic factors such as climate, land alteration, and pathogenic infection (Belsky and Joshi 2019). The environment is a major source of oxidative stress as it exposes insects to pathogens and pesticides (Gao et al. 2022).

Insects, specifically bees, are extremely vital to agriculture for their pollination services (Siviter et al. 2023). Pollinators in urbanized and agricultural landscapes are especially sensitive to chemical fumigants and habitat fragmentation (Quigley et al. 2019). Pollination is essential in agricultural settings, yet these settings are not often designed to support them (Hladik et al. 2016). Modern agriculture in the U.S. is dominated by conventional farms utilizing chemical methods of pest control and organic farms utilizing non-chemical methods (Shennan et al. 2017). When compared to conventional farming, organic farming is considered sustainable.

In the agricultural setting, pesticide use can negatively affect pollinator health (Brittain et al. 2010; Ma et al. 2019; Siviter et al. 2023; Wen et al. 2021). Organic practices aim to use pesticides developed from natural products as an ecological approach to preserve biodiversity (Baker et al. 2020). Insect abundance and plant diversity are typically higher on organic farms (Gabriel et al. 2013). This is likely related to the availability of nutrition on farms with more plant diversity (St. Clair et al. 2020). Other research also examine a third landscape termed ‘roadside’ as a least managed type (Nooten and Rehan 2019). These farms use minimum chemical pest management and thus represent the control condition.

Exposure to various pesticides, pathogens, and environmental conditions increases the production of ROS in honeybees (HB, *Apis mellifera*) (Olgun et al. 2020). The herbicide atrazine was related to elevated oxidative stress in honeybees (Williams 2016). Insecticides flupyradifurone and sulfoxaflor caused an increase in ROS, which could lead to oxidative stress (Chakrabarti et al. 2020). *Varroa destructor* is a parasitic mite that infects honeybee colonies significantly (Zheng et al. 2023). A study of honey bee pupae infested with *V. destructor* showed that compared to non-infested pupae, the infested had antioxidant levels two times higher (Badotra et al. 2013).

The accumulation of oxidative damage may contribute to senescence, or aging, in bees (Margotta et al. 2018). Additionally, several studies have demonstrated that honey bees may experience a tolerance to oxidative damage based on environmental or physiological factors (Li-Byarlay et al. 2016). Feral honey bees experience higher levels of OX as a tolerance mechanism than commercial honey bee colonies (Ward et al. 2022). Vitellogenin, a reproductive protein that has antioxidant properties, in female worker bees can lessen the effects of oxidative stress (Seehuus et al. 2006). Male honeybee drones manage oxidative stress by tolerating the lipid peroxidation it causes (Li-Byarlay et al. 2016). The bacteria within the digestive tract of honeybee queens have been found to undergo changes throughout their lifespans that may contribute to their longevity. Worker bees lack this potentially beneficial quality (Anderson et al. 2018). Co-infestations were more detrimental to honeybee health overall, although single infestations and infections of *Varroa* and DWV were related to lower total antioxidant status of bees (Łopieńska-Biernat et al. 2017).

Native pollinators are also essential to agriculture. Biodiversity of pollinators contributes to pollination success (Klein et al. 2007). Small carpenter bee (SCB, *Ceratina calcarata*) is a species native to North America (Nooten and Rehan 2022). However, research regarding stress mechanisms in SCB is not as extensive as research on honeybees. Wild bees are sensitive to ecological changes that occur around them (Harrison et al. 2019). When compared across landscapes of varying levels of management, the body size of *C. calcarata* offspring on intensely managed farms were smaller than those on less intensely managed farms (Nooten and Rehan 2019). The decline in body size observed during the experiment could indicate a decline in populations of wild bees as their offspring become less viable. Increases in temperature have a similar effect on body size, further demonstrating the relationship between feral bees and their environment (Kelemen and Rehan 2021). Another aspect of agricultural land use that has been detrimental to bees is grazing, which had a negative impact on the abundance of wild bees during a study in New Hampshire (Odanaka and Rehan 2019).

Even though pesticides are harmful to both managed and native bees, how they affect the oxidative stress in two bee species (*A. mellifera* and *C. calcarata*) in different farm landscapes is still unknown. The aims of our project were to investigate the changes of oxidative stress and pesticide exposures in different farm landscape (conventional, organic, and roadside as non-managed) in pollinator health. We test the hypothesis that (i) both pollinators experience high oxidative stress in conventional farm landscape, (ii) pollen from conventional farm landscapes will contain more pesticide residues than organice or roadsize landscapes.

## Methods and Materials

### Farm sites

Three farm landscape types were utilized in Central Ohio: organic farms, conventional farms, and roadsides. Sampling took place annually from 2021 to 2023, with three sites per landscape type (nine total). Site selection was based on farmer collaboration. While roadside sites remained consistent throughout the study, one organic and one conventional site were replaced after 2021 due to access limitations.

### Bees

At each farm site, two standard Langstroth honey bee hives (8-frame hives, 2 deep boxes) were installed in 2022 and 2023 from April to September in each farm. Each hive had eight frames containing 6000-7000 worker bees and a normal laying queen. All hives were started with same amount of honey and pollen supply at the beginning of the season. New pollen traps (Mann Lake, Hackensack, MN, USA) deployed for one week between mid-May and mid-June. For SCB, 100–120 assorted raspberry stems (45-50 cm long) tied with bamboo sticket (45-50 cm long) were placed at each site in late April as in other reports (Nooten and Rehan, 2019). Sticks were collected in June. Figure 1 shows the experimental design.

**Figure 1.**
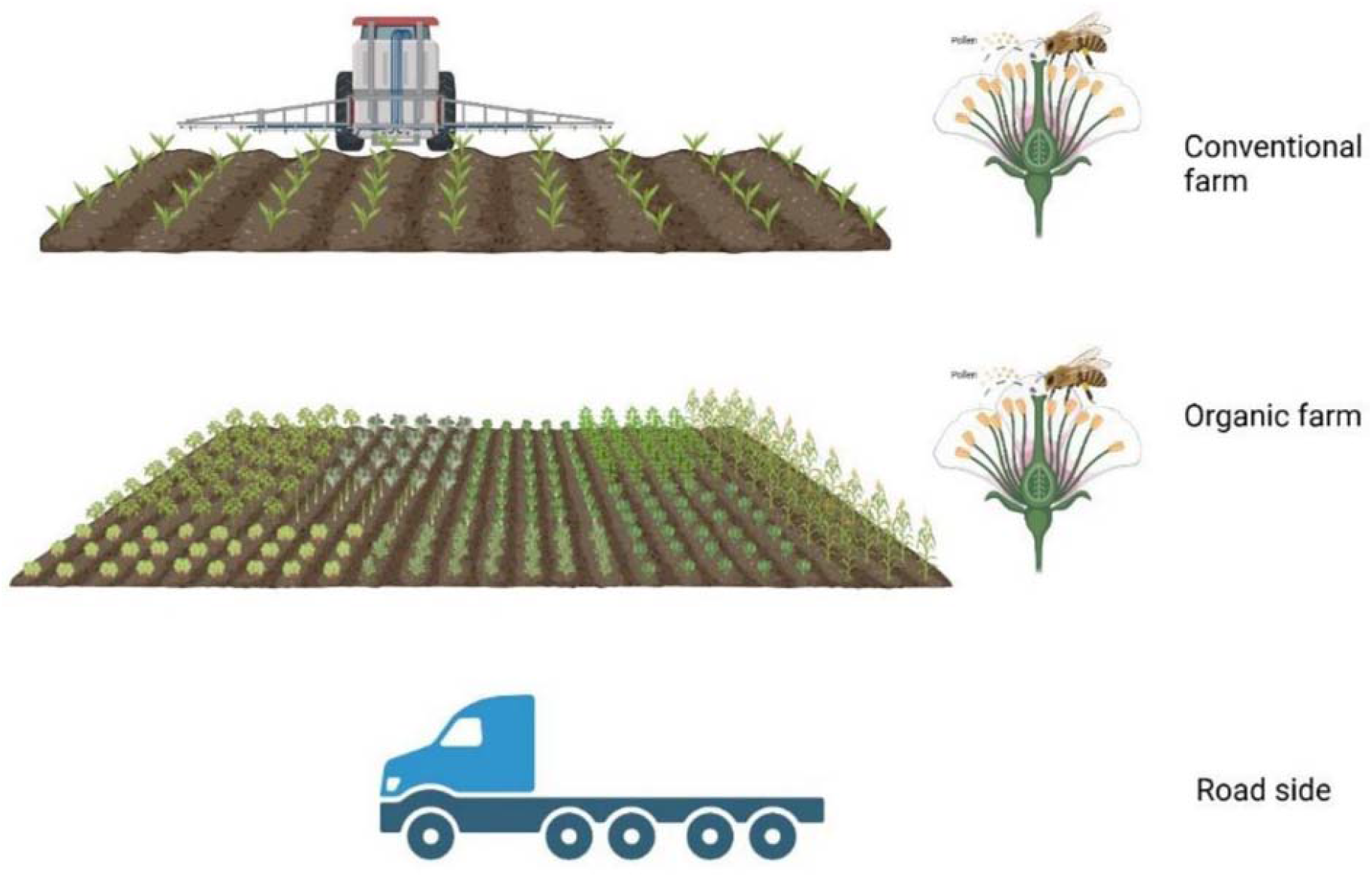

All *Ceratina* bees including adults and larvae were immediately collected and separated on the day we removed them from each field site. *Ceratina* species were identified using Discover Life (https://www.discoverlife.org/). Adults and larvae were flash-frozen in liquid nitrogen and stored at – 20°C. From May-June of 2021, 108 samples of adult HB and SCB were gathered from three types of landscapes (3 farms/type) (Figure 1, Created in BioRender. (2025) https://BioRender.com/w27f796). All samples were flash frozen by liquid nitrogen, and then stored in –80° C freezer until needed for experimentation.

### Tissue Preparation TBARS/BCA

One hundred-eight total adult and larval HB samples and twenty-seven adult SCB were prepared. HB sample preparation followed procedures outlined by Li-Byarlay et. al. 2016, Ward et. al. 2022. In brief, the whole head or larva of an individual HB was dipped in liquid nitrogen and homogenized in a 280µl solution of phosphate buffered saline 1X. For SCB, the entire bee was used. Supernatant was set aside for both assays. The solutions were then stored in -80 degrees freezer until needed for further processing. A combination of the Thio-barbituric Acid Reactive Substances (TBARS) and Bicinchoninic Acid (BCA) assays determined the level of oxidative stress. The TBARS (TCA Method) Assay kit (Cayman Chemical Company, Item catalog 700870, Ann Arbor, Michigan, USA) detects malondialdehyde (MDA), an indicator of lipid peroxidation. The BCA assay (Thermo Scientific, Rockford, Illinois, USA) measured total protein content. Readings were taken using the Synergy LX Multi-mode Plate Reader (BioTek, cat city state). The colorimetric procedure was followed for both assays. Measured protein content normalized the MDA concentrations, as explained by Li-Byarlay et. al. 2016 and Ward and Li-Byarlay, 2022.

### Tissue Preparation Protein Carbonyl HB and SCB

Protein carbonylation was determined using the OxiSelect Protein Carbonyl Fluorometric Assay kit (Cell Biolabs, Inc.). Samples preparation resembled methods of Li-Byarlay et. al. 2016, however 85 microliters of stock fluid was set aside for BCA. In brief, one-fourth of an adult honey bee thorax was homogenized in 200 microliters of Sample Diluent 1X following flash freezing by liquid nitrogen. Results from BCA were used to normalize carbonylation levels as suggested by assay protocol. Protocol was then followed exactly, excepting the adjustment of protein concentration for the tissue homogenate samples. Small carpenter bee adults from 2023 were also evaluated for protein carbonylation. The whole adult specimen was used. Samples preparation and evaluation is then the same as for adult honey bees.

### Pollen Samples and Pesticide Residue Analysis

Additionally, pesticide residue analysis of pollen samples from 2021 was conducted using GCMS (gas chromatography mass spectroscopy) and LCMS (liquid chromatography mass spectroscopy) at University of Guelph Agricultural Food Laboratory.

### Data analysis

Lipid peroxidation level is the MDA concentration in nanomoles over the total protein concentration in milligrams. Protein carbonylation is the protein carbonyl concentration in nanomoles over the total protein concentration in milligrams. For TBARS assay, an one-way ANOVA (analysis of variance) with post-hoc Tukey HSD (honestly significant difference) was conducted for both TBARS and protein carbonyl assays. Total sample size of hb is 54 bees, 6 adults or larvae were tested per farm, 9 farms total. One outlier was removed because it was out the range of mean ± 2 standard deviation. Twenty-seven adult SCB were compared for lipid peroxidation in the same manner.

Protein carbonyl levels were compared across fifty-six adult honey bees (n=56); Nine or ten honey bee thoraxes were used from each farm. Fifty-four adult small carpenter bees were tested for protein carbonylation (n=54). Recorded oxidative stress level is expressed in nmol/mg. Eighteen samples from each landscape were tested. Fifty-three were included in the analysis; One outlier was removed from the conventional group (n=17) (after applying IQR rule).

## Results

### OX and lipid damage in HB Adults

Average levels of lipid peroxidation in adult honeybees were as follows: 1.17, 1.67, and 0.96 nmol/mg for samples from organic (n = 18), roadside landscapes (n = 18) and conventional (n = 18), F _(2, 50)_ = 21.37, p-value < 0.001. Samples from the roadside landscape is significant higher than the organic landscape (p < 0.01, Tukey HSD Q statistic = 6.24), and conventional landscape (p < 0.01, Tukey HSD Q statistic = 9.01) (Figure 2).

**Figure 2.**
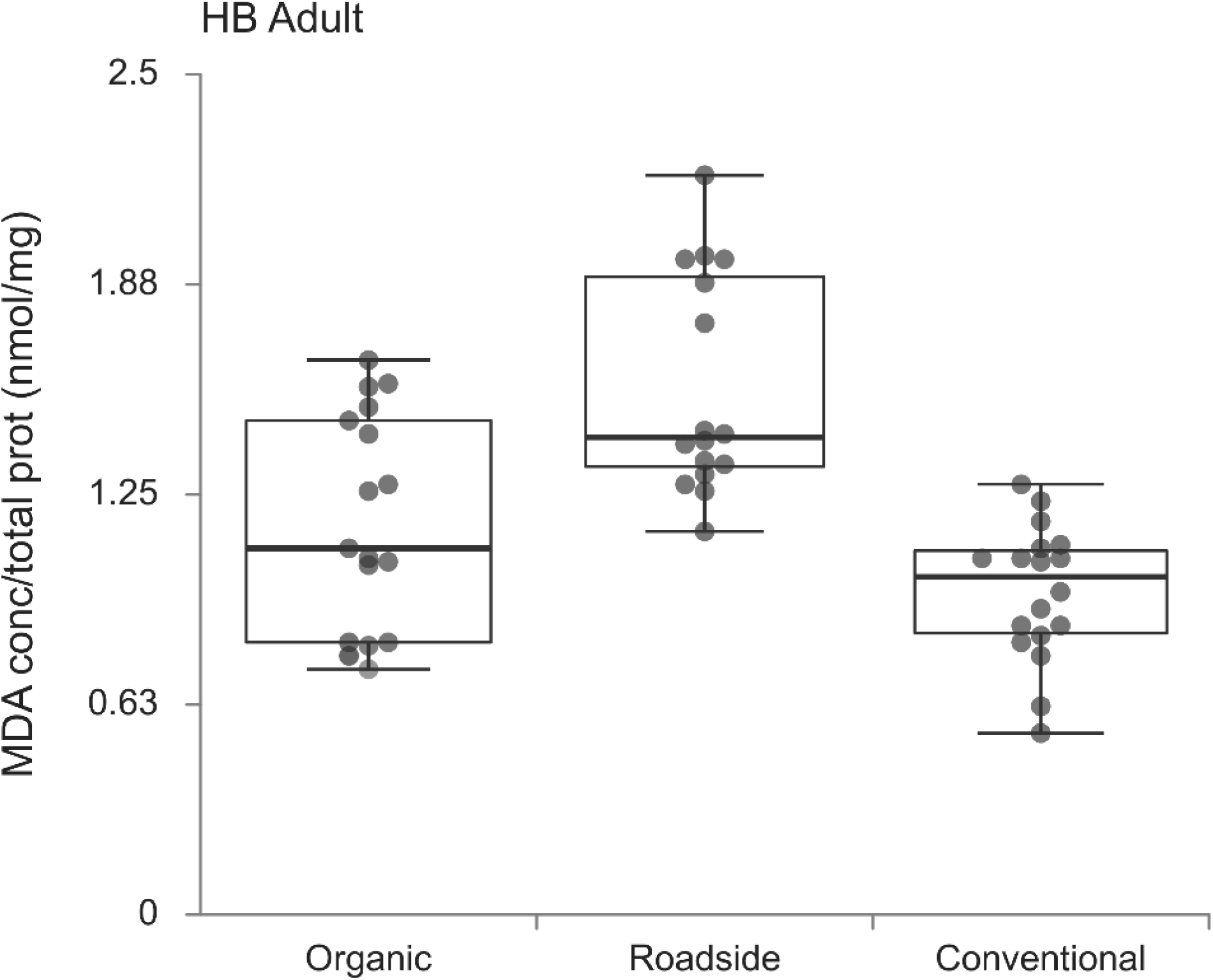

### OX and lipid damage in SCB Adults

Twenty-seven adult small carpenter bees were tested for lipid peroxidation (n= 27). The mean of each group is as follows: 1.20 nmol/mg for organic samples (n = 9), 1.84 nmol/mg for roadside samples (n = 9), and 1.65 nmol/mg for conventional samples (n = 9). F _(2, 24)_ = 3.65, p-value < 0.05. The level of lipid damage of samples from roadside landscape is significantly higher than those from the organic landscape (Tukey HSD Q statistic = 3.71, p < 0.05) (Figure 3). MDA levels in adult SCB were higher when compared to adult HB.

**Figure 3.**
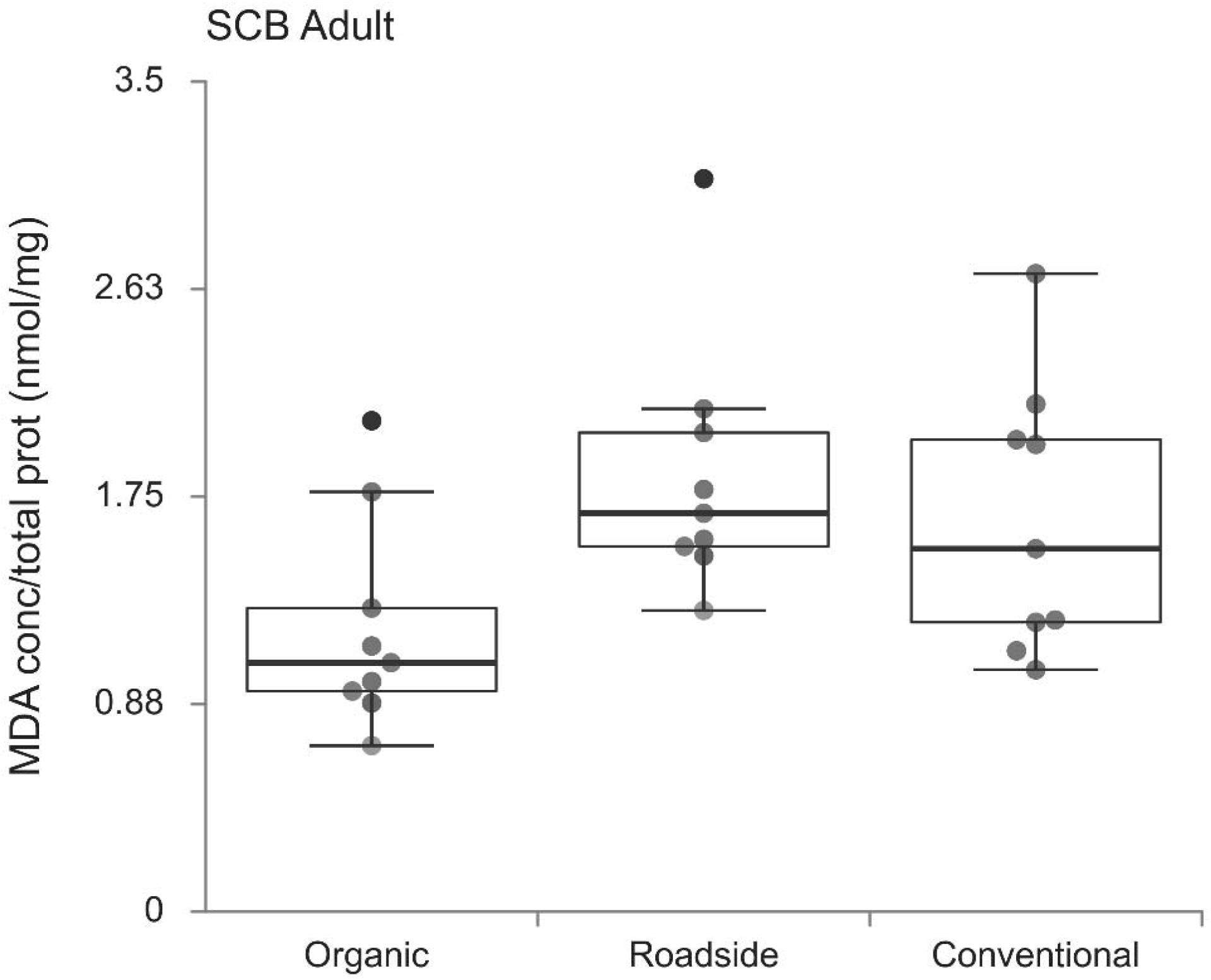

### OX and lipid damage in HB Larvae

The results of the analysis of larvae samples followed a similar trend to adult honey bees. Larvae from the roadside had the highest level (1.05 nmol/mg, n = 17), organic farms had the lowest levels of lipid damage (0.57 nmol/mg, n = 18). The average MDA level of samples from conventional farms was 0.85 nmol/mg (n = 17) (Figure 4). F _(2, 49)_ = 10.92, p-value = 0.0001. The level of lipid damage of samples from roadside landscape (Tukey HSD Q statistic = 6.57, p < 0.01) and conventional landscape (Tukey HSD Q statistic = 3.78, p < 0.05) are significantly higher than those of organic landscape.

**Fig 4.**
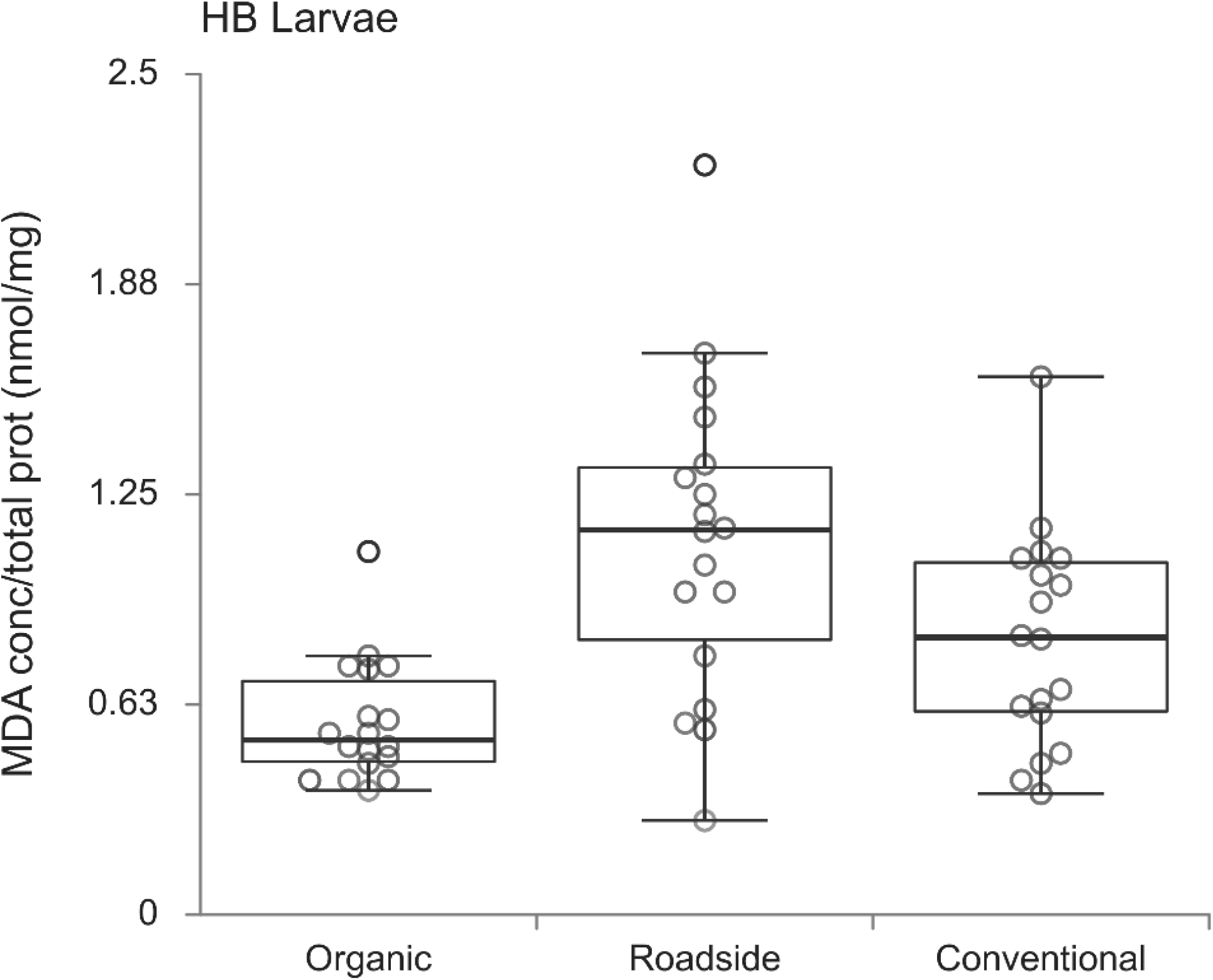

### Protein carbonylation in HB and SCB adults

Fifty-six adult honey bees (n = 56) were evaluated for oxidative stress included samples from organic landscapes (n = 18), the roadside landscape (n = 20) and conventional (n = 18). The means for 3 landscapes are as follows: 15.12, 18.48, and 20.49 nmol/mg for organic, roadside, and conventional landscapes (F_(2, 53)_ = 3.36, p value = 0.04). There is a significant difference between organic and conventional landscapes (Tukey HSD Q statistic = 3.62, p < 0.05) (Figure 5).

**Fig 5.**
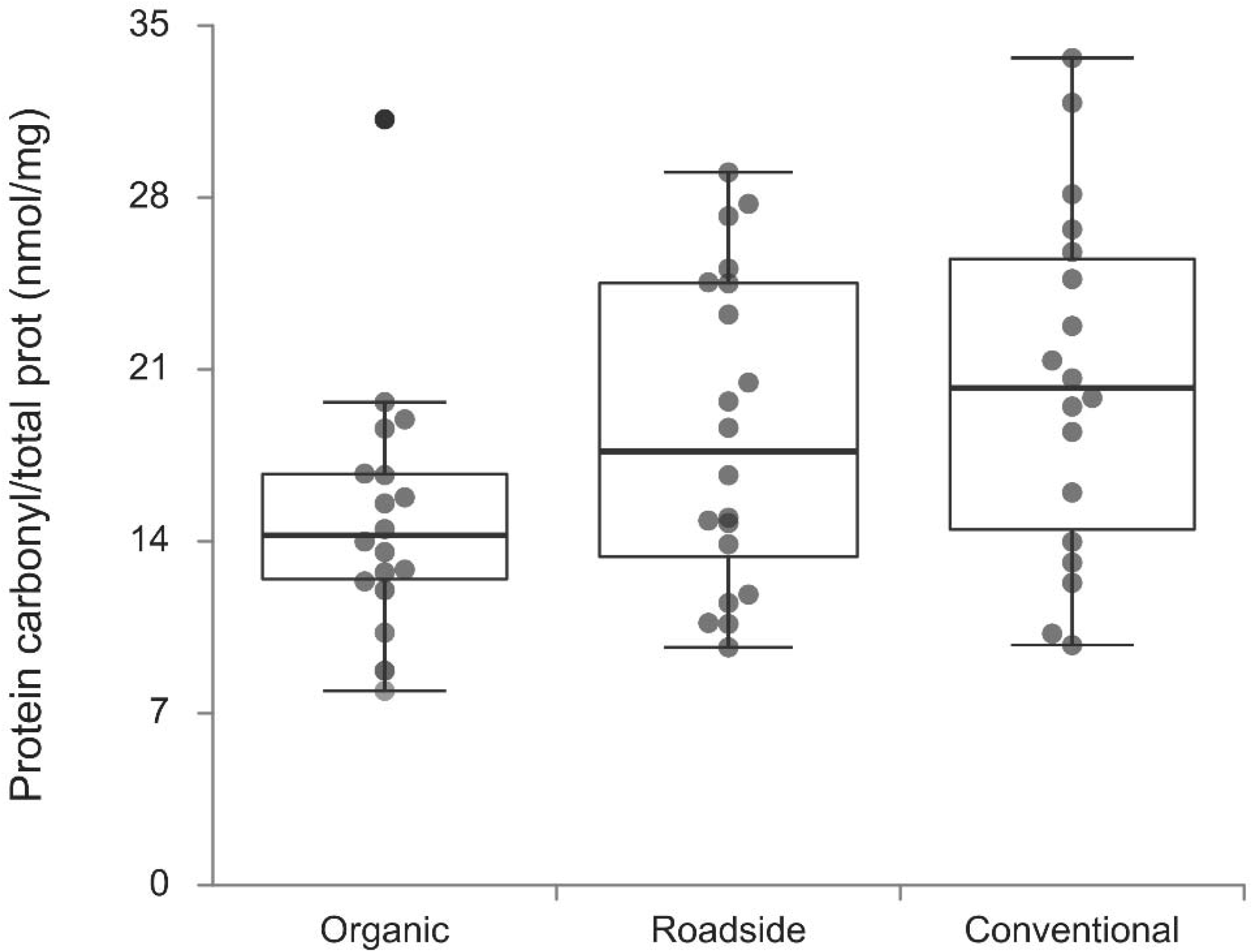

Twenty-six adult small carpenter bees (n = 27) were evaluated for protein carbonylation among three landscapes. The means for the levels of protein carbonylation of samples from 3 landscapes are as follows: 15.89, 14.58, and 14.74 nmol/mg for organic, roadside, and conventional landscapes (F_(2, 24)_ = 0.07, p value = 0.94). There is no significant difference among all landscapes (Figure 6).

**Fig 6.**
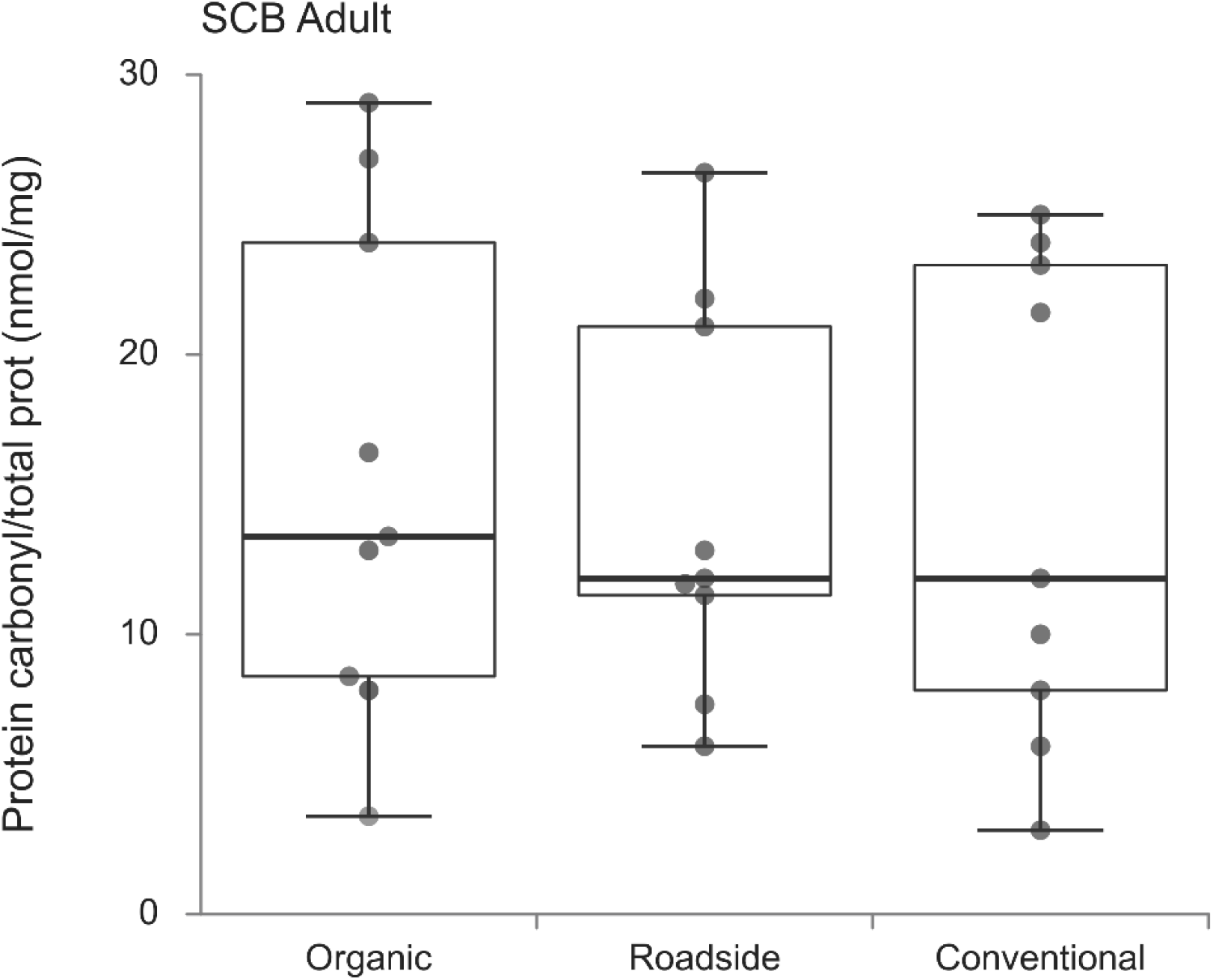
Comparison of the protein carbonylation levels in box plots among samples from 3 landscapes. P > 0.05 (nonsignificant). (B) Boxplots to show the levels of protein damage from each location (Fa, Fairborn; Ce, Cedarville; Be, Bellefontaine) as paired colonies (F, feral; M, managed).

### Pesticide Residue in Pollen

Results from residue analysis are expressed in parts per million (Table 3). Residue from conventional farms had the highest overall concentration of pesticides. Lower concentrations of pesticides were detected on roadside farms. However, in comparison to organic farms more types of pesticides were detected in conventional and organic pollen samples (Figure 7).

**Fig 7.**
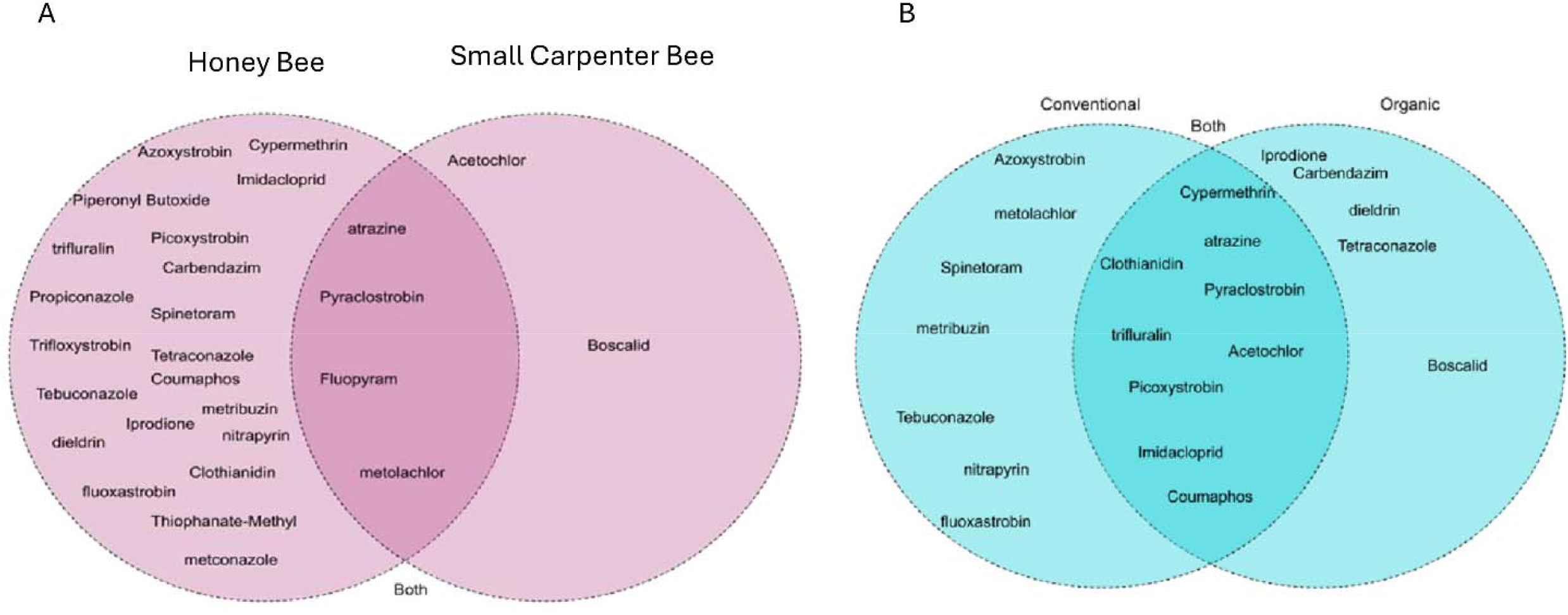

## Discussion

This study presents comparative data on oxidative stress in honey bees and small carpenter bees collected from three distinct landscapes, conventional, organic, and roadside. Our findings demonstrate a clear landscape-dependent variation in oxidative damage, with significantly elevated lipid peroxidation levels (MDA) in roadside-collected bees compared to those from organic or conventional landscapes.

The heightened MDA levels in roadside bees strongly suggest increased physiological stress, potentially due to chronic exposure to low-level pesticide residues and vehicular pollutants. This aligns with earlier work showing that sublethal pesticide exposure can disrupt detoxification pathways and induce oxidative damage in pollinators (Boncristiani et al., 2012; Christen et al., 2018). In contrast, bees from organic farms exhibited the lowest oxidative stress markers, consistent with lower or undetectable pesticide residues in those landscapes.

Interestingly, despite both species showing similar trends in oxidative damage across landscapes, HB displayed generally lower MDA levels than SCB. This may suggest species-specific resilience or differing foraging ranges and behaviors that influence exposure. Other research has highlighted that other wild bees such as bumble bees may forage over smaller, more diverse floral patches than honey bees, potentially reducing exposure to concentrated pollutants (Goulson et al., 2008).

The pesticide residue profiles we obtained also reveal that neonicotinoids and fungicides persist even in non-agricultural settings, although concentrations were markedly higher in roadside bees. This finding reinforces concerns that pollinator exposure to agrochemicals extends well beyond agricultural boundaries and may involve multiple stressors, including air pollution and habitat fragmentation.

## Conclusion

Our results provide clear evidence that landscape context influences oxidative stress levels in two key pollinator species. Elevated oxidative damage in bees from roadside habitats highlights the potential compounding effects of environmental pollutants and sublethal pesticide exposure. These physiological markers, combined with pesticide residue analysis, can serve as valuable tools for monitoring pollinator health across landscapes. Given the essential ecosystem services provided by bees, our findings underscore the urgent need for landscape-level strategies to reduce pollinator exposure to chemical stressors. Future studies should expand on these results by incorporating seasonal variation, behavioral data, and gene expression analyses of antioxidant and detoxification pathways.

## Acknowledgments

We thank Taya Barber, Joel Barhorst, Jairus Burrows, Austin Carey, Ashley Cordle, Zicheng Guo, Danielle Kroh, Ella Lowe, and Geremiah Rodgers for field assistance. KB and HLB are supported by the USDA grant award 2021-38821-34576 & NI241445XXXXG004.

## Author contributions

HL-B designed the experiment. KB, MLB, HL-B conducted field studies and collected samples. KB prepared samples, conducted data analysis, and wrote the first draft of the manuscript. RJ and SR provided resources. All authors reviewed the final manuscript.

## Data availability

Data will be provided when requested.

